# Characterizing Simultaneously Recorded Auditory Brainstem and Middle Latency Responses Using the Parallel Auditory Brainstem Response Paradigm

**DOI:** 10.1101/2025.09.11.675715

**Authors:** Isabel N. Herb, Melissa J. Polonenko

## Abstract

**Purpose:** The auditory brainstem response (ABR) and middle latency response (MLR) are used to characterize hearing but typically require separate recordings due to differing optimal parameters and setups in clinical systems. Advances such as the parallel ABR (pABR) paradigm now allow simultaneous recording of both responses, enabling more comprehensive assessments. We evaluated this simultaneous recording approach by characterizing responses at near- and suprathreshold levels using parameters optimized for pABR.

**Methods:** This study used an open dataset [1] of pABR recordings from 20 normal hearing adults to tone-pip stimuli at five frequencies, six presentation rates, and two intensities. ABR (I, III, and V) and MLR (P0, Na, Pa, Nb, Pb) waves were analyzed for presence, latency, and amplitude using linear mixed effect modeling.

**Results:** At both levels, ABR wave V and MLR waves Na and Pa were the most consistently identifiable and largest peaks across frequencies, rates, and participants. Frequency and rate changes affected ABR wave V amplitude more than the MLR peaks, whose amplitudes and latencies remained comparatively stable across the moderate and soft levels used in this study.

**Conclusion:** ABR wave V and MLR waves Na and Pa are robust and consistently identifiable with pABR-optimized parameters. A simultaneous recording approach that leverages visualization of all three components could both assess auditory function across more areas of the auditory system using a moderate suprathreshold level, and support detection/interpretation of smaller wave Vs for threshold estimation, particularly for broader low frequency responses.

## Introduction

Auditory evoked potentials are used to characterize hearing function across the loudness range, from detecting sound to processing sound at high levels. The most widely used evoked potential in the clinic is the auditory brainstem response (ABR), which reflects transmission of sound from the auditory nerve to rostral brainstem [2–5]. The middle latency response (MLR), although less studied, also provides valuable information about hearing function from the rostral brainstem through the thalamus and auditory cortex [6–9]. Therefore, both responses are important for a more comprehensive neural assessment of hearing. However, each response must be separately recorded in current clinical systems because of differences in analysis time windows required by the periodic stimulation paradigms. But time constraints during a clinical appointment often render only one type of recording possible to complete, which is usually the ABR for its reliability and well-established role in assessing auditory function and predicting hearing threshold [9–12]. However, simultaneous recordings are now possible with recent advances in recording paradigms that use randomized instead of periodic stimulation, thereby allowing extended analysis windows [13–15]. But simultaneous recording requires the same parameters be used for both responses, which may be optimal for one (ABR or MLR) but suboptimal for the other. This paper explores the feasibility of combined ABR and MLR assessment using ABR-optimized parameters of one such paradigm, the parallel ABR [13, 14].

The pABR paradigm was created to reduce testing time for ABR exams that estimate hearing thresholds, but confers the added benefit of extended analysis windows to view the MLR without modifications to the recording protocol [13, 14]. The extra information available from simultaneously viewing both responses may support more accurate or confident threshold estimation than when using the ABR alone as well as provide a more complete evaluation of auditory function throughout more areas of the auditory system [9, 13, 14, 16]. But whereas wave V has been characterized for tone-pip ABRs in general [11, 12, 17–19] and when measured with the pABR paradigm [13, 14], little research exists on MLR responses to tone-pips and especially when stimuli are presented at softer levels nearer hearing threshold [7, 9, 20–23]. Therefore, the goal of this study was to characterize MLR waveform components at two levels – a moderate level to assess suprathreshold function and a level close to threshold (so that responses are still present) – to determine whether simultaneously collecting MLR with the ABR is feasible without changing the ABR protocol.

### Overview of ABR and MLR Responses

ABR and MLR responses are scalp-recorded potentials generated in response to auditory stimuli and reflect activity from the level of the auditory nerve to auditory cortex. The ABR consists of five wave components (waves I–V) that occur within the first 10-15 ms after stimulus onset. Each wave is associated with a neural generator located in specific regions of the auditory pathway. The three most prominent waves – waves I, III, and V – are generated in the distal portion of the auditory nerve, cochlear nucleus, and from projections between the lateral lemniscus and inferior colliculus, respectively [2–6]. The MLR follows the ABR in time and consists of a series of alternating positive (P) and negative waves (N): P0 (∼10-15 ms), Na (∼15-22 ms), Pa (∼24-24 ms), Nb (∼35-50 ms), and Pb (∼50-60 ms) [24]. MLR components are generally attributed to the thalamocortical pathway, with Na arising from the midbrain, medial geniculate body, and thalamus, and Pa, Nb, and Pb from successive regions of the auditory cortex [6, 7, 24, 25]. Thus, the MLR provides insight into regions of the auditory system rostral to the ABR generators, reflecting higher level auditory processing [6, 7, 9, 16, 20, 22, 23].

Beyond their neural generators, the ABR and MLR also differ in their standard recording parameters. First, MLR protocols use lower stimulation rates than ABR protocols. The longer duration between periodic presentations of the stimulus allows for longer post-stimulus analysis windows that capture the entire MLR waveform.

Furthermore, the slower rate evokes larger amplitudes by allowing more synchronized firing from the thalamocortical generators that have slower temporal properties [6, 22, 26–28], as compared to the early structures generating the ABR that can quickly recover and respond to high presentation rates [15, 26–28]. Studies generally report a sharp decrease in MLR amplitude with an increasing stimulation rate to about 40 Hz, but minimal latency changes [22, 26–28]. Second, MLR filtering parameters are lower (10-500 Hz) than those for ABR (30-2000 Hz) to match the spectral characteristics of each response [6, 12, 22, 26–28]. Third, patient state/arousal can influence the presence and morphology of the MLR, particularly during early developmental periods. While often still recordable and visible, MLR amplitudes can decrease during quiet sleep [8, 27, 29], which may make responses more variable or difficult to interpret. For this reason, MLRs were not routinely adopted into pediatric clinical settings in favor of using the ABR, where testing is done during sleep to minimize noise and movement to collect viable responses. But simultaneous recording may make use of the extra information provided by the MLR, when present, to support clinical interpretations.

Few studies have explored whether modern recording systems can reliably measure the ABR and MLR together. While the optimal parameters for each response traditionally differ, both responses could be measured simultaneously using a shared set of recording parameters that, while possibly suboptimal for one or both responses, preserve key response features that provide useful information. While the ABR remains the primary clinical measure, reliably observing the MLR in the same recording could justify its concurrent use in clinical protocols. We address the feasibility of simultaneous recording by evaluating the MLR presence when using optimal parameters for the ABR. Recording parameters that can reliably capture both responses paves the way for more comprehensive assessments of auditory function without extending the appointment duration.

### ABR and MLR for threshold estimation

Tone-pip ABRs are mainly used to estimate hearing thresholds in individuals who cannot provide behavioral responses. Wave V is more reliably identified and correlates better with behavioral thresholds than MLR, making it the gold standard for estimating thresholds in infants [8, 27, 29–31]. While MLRs can be used to estimate hearing thresholds [9], they are less accurate for infants and children due to the greater effect of arousal on MLR amplitudes during central auditory development [8, 27, 29]. Pa develops first, becoming adult-like around 10 years of age, while Pb becomes adult-like last, at around 15 years of age [8, 24, 29]. Due to the greater variability in infant MLR responses during sleep, MLR-based threshold estimation was largely abandoned, especially for systems with limited time windowing that forced clinicians to choose either the ABR or MLR paradigm.

Although arousal state is an important consideration, the MLR is often still present during light and rapid eye movement (REM) sleep and only begins to diminish as the individual enters deeper sleep stages [8, 9, 27, 29]. Therefore, although it may not be the most reliable standalone method to detect threshold, recording the MLR simultaneously with the ABR may enhance threshold detection when present. Compared to the ABR, MLR components tend to be larger at lower frequencies, and less sensitive to stimulus level changes [6, 8, 9] because they reflect cortical activity that integrates slower temporal information over a longer time window, allowing for better synchronization with low frequency stimuli. This contrasts with the ABR, where low-frequency stimuli often produce smaller, later, and broader responses that are closer to the background noise floor at low intensities, making wave V detection and threshold estimation more difficult [3, 10, 11, 17, 23]. Therefore, an optimal recording paradigm would leverage the benefits of larger response amplitude for the ABR at high frequencies and for the MLR at low frequencies.

### Suprathreshold measures offer insight to auditory function beyond detection

Suprathreshold auditory evoked potential testing uses high-intensity stimuli to evoke responses with prominent component waves. Comparing the amplitudes and latencies of these waves, as well as the relative timing between the waves, assesses function at different points in the auditory pathway. This method was used to help identify the likely presence of an acoustic neuroma/vestibular schwannoma before MRI proved to be more sensitive for accurately detecting smaller tumors [5, 11, 16, 25]. Today, clinical suprathreshold testing mainly uses high-intensity clicks to diagnose auditory neuropathy spectrum disorder [ANSD; 10, 11, 16, 30, 31]. In ANSD, ABRs are characterized by preserved early cochlear potentials (the cochlear microphonic) but absent or poorly formed ABR waves [10, 11, 16]. If present, however, MLRs could indicate partial preservation of synchrony (or reduced disruption of synchrony) along the thalamocortical pathway [7, 8, 16, 24].

Current research also uses suprathreshold auditory potentials to investigate auditory processing and neural function in individuals who report difficulty hearing in noise despite having normal audiometric thresholds [32–36]. These suprathreshold methods are often conducted at very high levels (70-100 dB nHL, or ∼90-120 dB peSPL) using clicks to elicit large ABR wave I responses. However, these paradigms have had mixed success in predicting behavioral challenges [32–36]. High intensity levels can also be distressing or intolerable for some groups, limiting clinical applicability [16, 32, 33, 35, 36]. Moreover, cochlear mechanics vary with level due to the nonlinear nature of the auditory system. The cochlea amplifies low-level sounds more than high-level sounds and the outer hair cells lose their amplification ability at higher levels, leading to a compressed response curve [32, 33, 35, 37]. As a result, responses recorded at high intensities may not reflect auditory processing under more typical listening conditions. Rather than using these high intensity levels, several studies have emphasized the importance of testing at suprathreshold levels closer to those encountered in daily listening [32, 33, 35, 36]. Given the importance of ecological validity, adding suprathreshold levels of moderate intensity to clinical protocols is worth considering. In this study, we analyzed suprathreshold responses at a moderately presented level (81 dB peSPL, or ∼60 dB nHL) that is lower than typically used in clinical settings. This allowed for the evaluation of whether a more ecologically valid level could still yield meaningful diagnostic information with the ABR and MLR.

### Simultaneous recording of ABR and MLR with one set of parameters

The pABR system’s ability to record responses simultaneously creates an opportunity to reintroduce the MLR into clinical and research testing. The key question that remains is whether useful MLRs are measurable when recorded with parameters optimized for measuring ABRs. Before suggesting a change towards a simultaneous ABR and MLR protocol, the feasibility of recording MLRs must be determined when using parameters optimized for the ABR. While Polonenko & Maddox [14] showed visible MLRs at faster stimulation rates, they focused on characterizing wave V. This study used their publicly available dataset [1] to characterize more components of the ABR and MLR obtained from the same recording using faster pABR rates. ABR and MLR responses at both near-threshold and suprathreshold levels were analyzed for their visibility and potential for use in clinical settings. At suprathreshold levels, the presence, amplitude, and latency of ABR and MLR components were evaluated. At lower levels, the detectability of Na–Pa was examined to determine whether the MLR could serve as a complementary marker for threshold estimation alongside wave V. These analyses assess the feasibility of simultaneous ABR and MLR acquisition using a single recording paradigm, with implications for more efficient and ecologically valid clinical protocols.

## Methods

### Dataset

Responses to pABR stimuli were obtained from an open repository on Dryad [1]. Details about stimulus and recording methods are described in the associated paper by Polonenko & Maddox [14]. In brief, the dataset includes 2-channel electroencephalography (EEG) recordings from 20 adults (13 females, 6 males, and 1 non-identifying participant) with normal hearing and a mean ± SD age of 22.5 ± 4.2 years (range: 18–35 years). Normal hearing was defined as ≤ 20 dB HL at octave frequencies from 250 to 8000 Hz. The pABR stimuli consisted of 500, 1000, 2000, 4000, and 8000 Hz tone-pips presented dichotically at stimulation rates of 20, 40, 60, 80, 100, and 120 Hz and intensities of 51 and 81 dB peSPL (∼33 and 63 dB nHL respectively, depending on frequency, see Polonenko & Maddox [14]). Each trial lasted 1 second. All 12 conditions (6 rates × 2 intensities) were interleaved and recorded for 10 minutes each during a 2-hour session. EEG signals were sampled at 10 kHz using two EP-Preamplifiers and a BrainVision actiCHamp amplifier (Brain Products GmbH, Gilching, Germany). Stimuli were presented at 48 kHz via an RME Babyface Pro soundcard (RME Audio, Haimhausen, Germany) and delivered through Etymotic Research ER-2 insert earphones. Digital triggers synchronized stimulus presentation and EEG recordings using a custom trigger box [38].

### pABR Derivation

Details of the pABR waveform derivation are provided in Polonenko & Maddox [13, 14]. Briefly, triggers marked the onset of each 1-second epoch, and EEG data from 500 ms before to 500 ms after each epoch were extracted to form a 2-second segment. Responses for each tone-pip were calculated using circular cross-correlation in the frequency domain between the EEG segment and the corresponding rectified impulse train that was used to create the pABR stimuli. The middle second was discarded and the remaining intervals were concatenated to yield the final response from −500 to 500 ms relative to the stimulus onset. To compute the average for each condition, each epoch was weighted by the inverse of its pre-stimulus baseline variance, minimizing the influence of noisy trials in a manner similar to Bayesian weighting [39]. Responses were then averaged across both EEG channels to further improve signal-to-noise ratio (SNR).

### Waveform Analysis

EEG data were filtered offline between 30–2000 Hz for ABR and 10–500 Hz for MLR responses using a causal first-order Butterworth bandpass filter, and with notch filters at odd multiples of 60 Hz to remove electrical noise. The MLR bandpass filter settings were determined by first comparing several ranges used previously in the literature [6, 7, 25–27, 29]. Of the most typically used filter settings (1–300 Hz, 10–500 Hz, 30–2000 Hz), the 10– 500 Hz range provided the clearest MLR components without latency changes relative to the ABR filter settings. While simultaneous evaluation of ABR and MLR with the same filter settings would occur in real-time, the lower range provided better conditions for selecting peaks when characterizing the broader MLR components.

Waveforms were manually inspected for the presence of ABR (I, III, V) and MLR (P0, Na, Pa, Nb, Pb) components. If present, peak amplitude and latency were manually marked. Grand average waveforms across participants for each condition served as templates to guide peak selection, particularly for broader peaks (e.g., ABR peaks for 500 Hz stimuli and MLR components Nb and Pb). Fig. 1 gives an example of the grand average template for the response to a 1000 Hz tone-pip, with all ABR and MLR components marked. Amplitude was measured from peak to following trough for positive components and from trough to following peak for negative components. For the MLR, this meant that each peak amplitude represented the difference in amplitude between subsequent components since there were no troughs between components like there is for the ABR. For example, Na amplitude was calculated from Na to Pa, Pa amplitude was calculated from Pa to Nb, etc. The peak and trough were chosen at the centermost position for broad waves. Responses from the opposite ear or other tone-pip frequencies were referenced to improve peak identification and selection of the where to mark the peak. Peak choices were also guided by replication responses derived from odd and even trials.

**Fig. 1.**
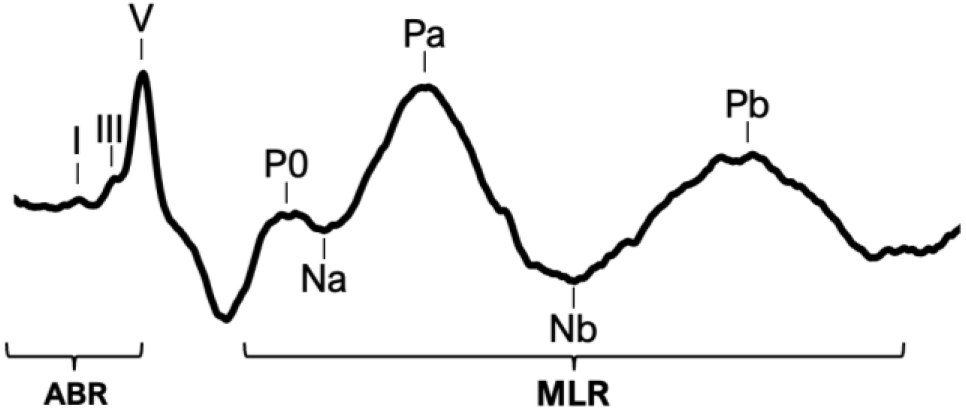
81 dB peSPL grand average waveform across participants, ears, and rate for the 1000 Hz tone-pip

Peaks were selected by both authors, who are trained audiologists. The first author picked peaks for all participants and the second author picked peaks from a random subset of 6 participant waveforms (30%) to evaluate inter-rater reliability [13, 14]. Inter-rater ICC3 values between the two authors were excellent (>0.90) or good (>0.75) for all peaks and levels, except for wave I and Pb latencies at 51 dB peSPL, which showed moderate agreement (0.74 and 0.64 respectively) [40]. This was expected, given wave I was small and often not present (see later section), and Pb was the latest, smallest, and broadest peak of the MLR. Therefore, these peaks were not included in the subsequent statistical analyses at the lower level.

### Statistical Analysis

Linear mixed-effects models were used to evaluate changes in ABR and MLR peak latencies and amplitudes across increasing stimulation rates at 81 dB peSPL. Frequency was input to the models as kHz. The data were log-log transformed for linearity: latency, amplitude, rate and frequency were transformed using log10. Because of the non-linear relationship, 19 was subtracted before logging rate so that the intercept of the log-log model would represent the values for our lowest measured rate (20 Hz). Five increasingly complex models were constructed and compared to determine the model that best explained the data. Model 1 was an intercept-only model. Model 2 added the fixed effects of rate, frequency, and peak. Model 3 added all 2-way interactions and the three-way interaction between rate, frequency, and peak. Model 4 added random effect intercepts for rate, frequency, peak, and ear by participant. Model 5 contained all effects from Model 4 but excluded the three-way interaction between rate, frequency, and peak. Stepwise elimination of Model 4 retained all random effects, so no further models were created.

The five models were compared using the corrected Akaike Information Criterion (AICc) [41], which adjusts for small sample sizes. Change in AICc (ΔAICc) values reflect how much worse each model is relative to the best model, and AICc weights indicate the relative likelihood that a given model is among the best in a set. As such, the values for AICc, ΔAICc, and AICc weights were compared across models to identify the model with the strongest empirical support. Comparative analysis of the models is given in Online Resource 1. After comparison, Model 4 gave the most support for both amplitude and latency. However, inspection of the full dataset revealed a notably divergent pattern for 8000 Hz amplitudes, which could be contributing to the complex model with 3-way interactions. To determine if a more parsimonious model explained the main amplitude trends for the other 4 tone-pips, amplitude models were re-analyzed excluding 8000 Hz. Model 5, without the three-way interaction, was the most parsimonious for amplitude (i.e., lowest ΔAICc). Model 5 also had a small ΔAICc of 5.42 [41] relative to Model 4 for latency (which included 8000 Hz for the latency models), supporting the simpler model’s use for both measures. Consequently, the model excluding the three-way interaction (Model 5) was selected to highlight the overall parameter effects, and is as follows:

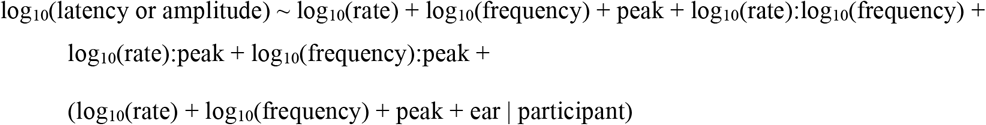

Ratios were used to evaluate the relative changes in peak metrics for the two stimulation levels. Amplitude and latency ratios were calculated for responses at 51 dB peSPL by dividing each metric by its corresponding value at 81 dB peSPL. Thus, an amplitude ratio of 0.5 indicates the amplitude halved, and a latency ratio of 2 indicates the latency was delayed by two times for the lower compared to higher level. Analysis focused on waves V, Na (Na– Pa), and Nb (Nb–Pb), based on their detectability (see Results section). Peaks I, III, and P0 were excluded due to their variable presence at the lower level. Only the 40 Hz stimulation rate was included as the most optimal single rate for pABR collection [13, 14], although data for all rates are provided in Online Resource 2. Outliers (n = 470, 4.8%) were excluded if they fell outside 1.5 times the interquartile range.

The ratios were evaluated in two ways. To determine if each peak significantly changed from the higher to lower level, independent two-tailed t-tests (µ = 1) were performed on amplitude and latency ratios for each peak-frequency combination, and *p*-values were corrected for family-wise error using false discovery rate [42]. To assess whether wave V amplitude and latency changed to a greater extent relative to Na-Pa and Nb-Pb, a series of increasingly complex linear mixed-effects models were fit, as done for the analysis with the higher level data: (1) random intercept for participant; (2) added fixed effects of peak and frequency; (3) added their interaction; (4) added random intercepts for ear and peak–frequency interaction; and (5) a final model selected via stepwise elimination for the random effects. Model comparisons using AICc (see Online Resource 3) revealed that the most optimal model(s) were the stepwise selected model of ratio ∼ peak + (peak | participant) for amplitude and ratio ∼ peak + log10(frequency) + (log10(frequency) | participant) for latency. However, the ΔAICc were small (2.43 for amplitude and 1.86 for latency) between these two stepwise models and the corresponding models that only included fixed effects plus a random intercept by participant (Model 2). Therefore, the latter, more parsimonious model was selected for both metrics, as described below:

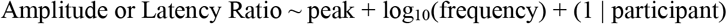

Statistical analyses were performed in RStudio [43] using R version 5.1 [44]. ICC3 values were calculated with the *psych* package [45]. Mixed-effect models were implemented using *lme4* [46], with analysis support from *broom*.*mixed* [47], and *lmerTest* [48]. T-tests were performed using the *stats* package [44]. Data wrangling and visualization were performed using *tidyverse* [49]. Waveform derivation and peak selection were completed using custom scripts implemented in Python and the use of *mne* [50]. Analysis code with documentation will be made publicly available on GitHub at https://github.com/polonenkolab/pabr_simultaneous_abrmlr.

## Results

### ABR and MLR components are measurable at a moderate suprathreshold level

At 81 dB peSPL, recordings were analyzed across all stimulation rates (20–120 Hz) to more fully characterize the different component peaks of the ABR and MLR. Fig. 2 shows the grand average waveforms for each ear, frequency, and stimulation rate. Overall, ABR peak morphology was sharper for higher-frequency tone-pips, whereas MLR peaks were more clearly defined for low-frequency tone-pips. These peaks were then quantified further for the presence, amplitude, and latency of ABR waves I, III, V and MLR waves P0, Na, Pa, Nb, and Pb, as shown in Fig. 3.

**Fig. 2.**
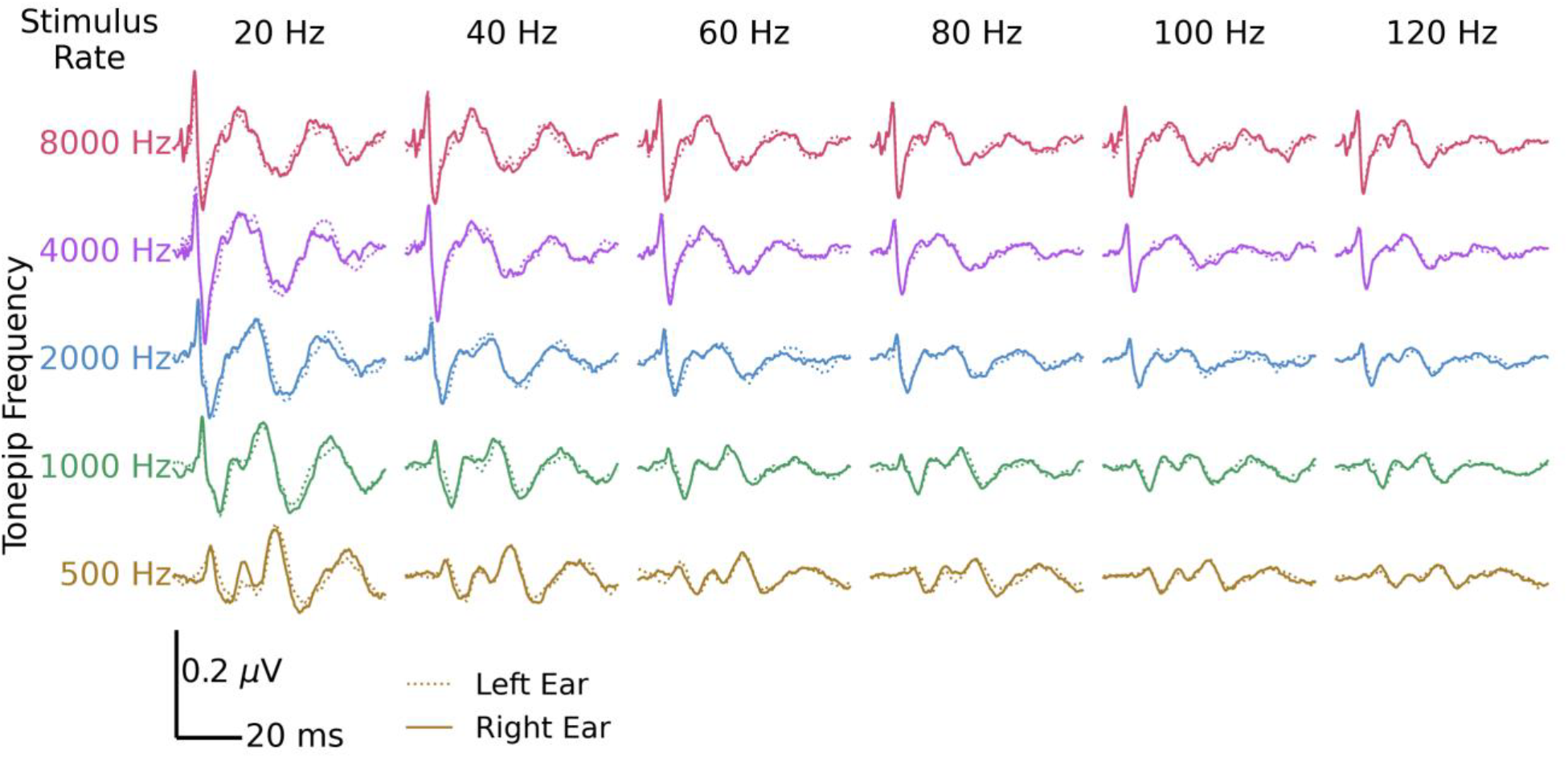
Grand average responses for 20 participants at 81 dB peSPL for each tone-pip frequency (rows) and rate (columns), separated by right (solid) and left (dotted) ears

**Fig. 3.**
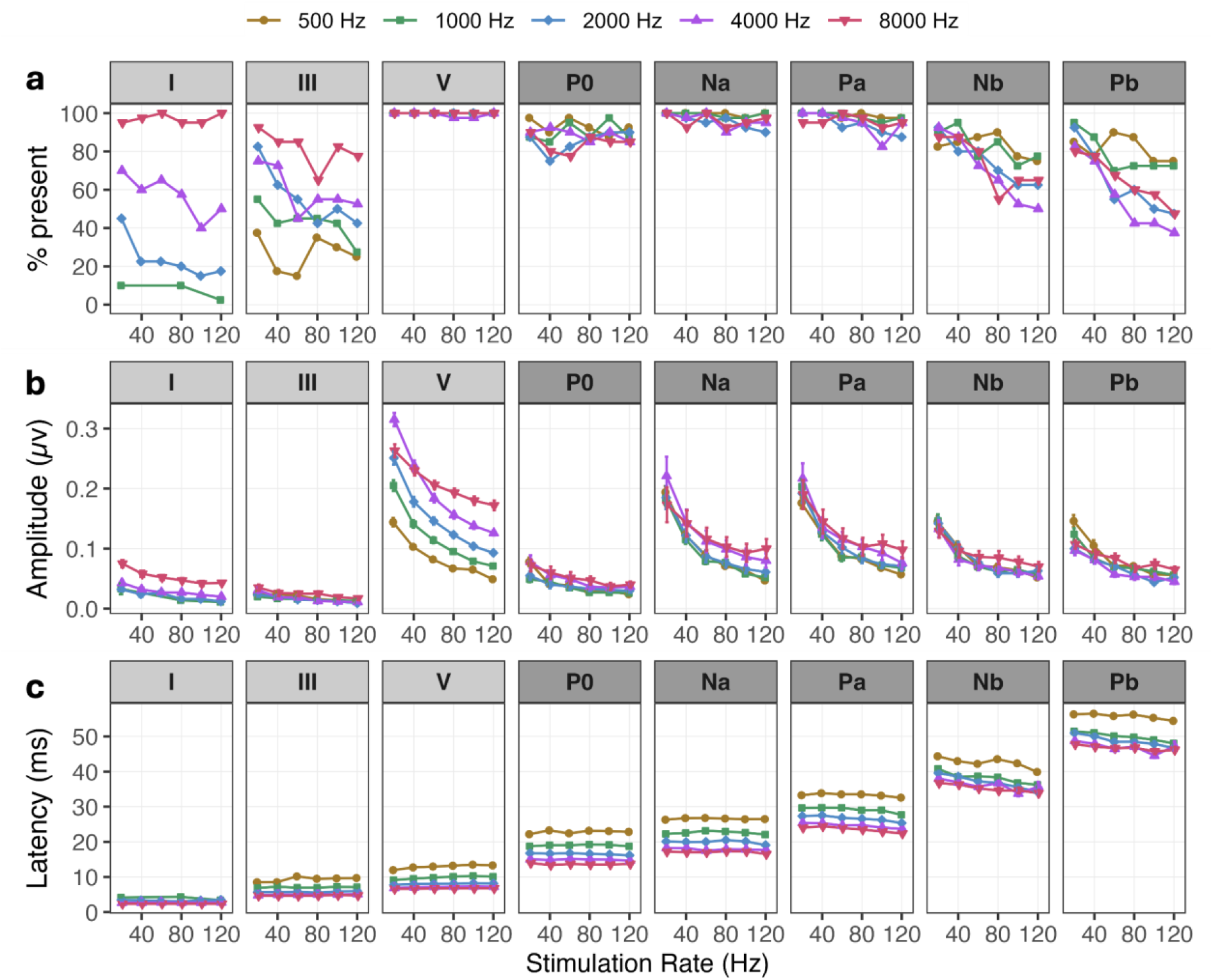
Wave (a) presence across participant ears, as well as the mean (b) amplitude and (c) latency for stimuli presented at 81 dB peSPL across 6 rates and 5 tone-pip frequencies. Colors and shapes denote tone-pip frequency. Error bars (where large enough to be seen) indicate standard error of the mean. Panel shading delineates ABR (light gray) from MLR (dark gray) waves

### ABR Wave V and MLR Waves Na and Pa are the most consistently present components across frequency and rate

Fig. 3a illustrates the proportion of identifiable waves across ears of participants at each frequency and stimulation rate. At this moderate level, the clearest and most consistently present waves across conditions were the ABR wave V and MLR waves Na and Pa. Each of these MLR waves were nearly 100% identifiable at the lowest rates (20-80 Hz), with a slight reduction at the higher rates (100 & 120 Hz) to 94.8% and 89.5%, respectively. In contrast, MLR components P0, Nb, and Pb were present on average in 79%, 80% and 84% of participants across conditions, and were most identifiable when using low frequency tone-pips presented at lower stimulation rates. Lastly, early ABR waves I and III were the least identifiable peaks: only high-frequency (≥ 4000 Hz) and low-rate (<60 Hz) stimuli evoked identifiable peaks in at least 80% of participants but were otherwise absent in at least half the participants (ranging from 0-55%).

### Frequency and rate parameters affect MLR peaks differently than ABR wave V

As shown in Fig. 3b, response amplitudes decreased non-linearly with increasing stimulation rate. ABR wave III showed the smallest amplitude overall, followed by wave I and the MLR component P0—peaks that were also the least frequently observed across participants (Fig. 3a), likely due to difficulty in distinguishing their small amplitudes from baseline noise. Due to the limited number of observations, waves I, III, and P0 were excluded from further analysis. All subsequent comparisons focused on ABR wave V and MLR components Na, Pa, Nb, and Pb. Furthermore, the 8000 Hz tone-pip response displayed a notably smaller increase in amplitude with decreasing rate than the other frequencies for every peak, resulting in an amplitude that was lower than the amplitude for 4000 Hz at the lowest rate (20 Hz). This unique pattern created a significantly more complex model that did not converge. Exclusion of 8000 Hz data confirmed that a simpler model without 3-way interactions best explained the main effects of rate and frequency for 500–4000 Hz tone-pips, which are also the most commonly used tone-pips in the clinic. In favor of clearer and simpler explanations of how MLR patterns compare to those of wave V, we present the simplest model that excludes 8000 Hz (see Methods for details). MLR wave Na was used as the reference in the model, which is provided in Table 1.

**Table 1.** Linear mixed effect model results for 81 dB peSPL.

|  | Amplitude (excludes 8 kHz) |  |  |  | Latency (includes 8 kHz) |  |  |  |
| --- | --- | --- | --- | --- | --- | --- | --- | --- |
|  | *Estimate ± SE* | *df* | *t* | *p* | *Estimate ± SE* | *df* | *t* | *p* |
| Intercept (Na) | -0.719 ± 0.035 | 21.70 | -20.73 | < .**.001** | 1.354 ± 0.007 | 23.07 | 206.31 | < .**.001** |
| Rate | -0.260 ± 0.007 | 246.80 | -36.84 | < .**.001** | 0.002 ± 0.002 | 125.15 | 1.18 | .240 |
| Frequency | -0.027 ± 0.036 | 32.54 | -0.76 | .451 | -0.154 ± 0.005 | 38.60 | -28.09 | < .**.001** |
| Pa | 0.004 ± 0.017 | 119.27 | 0.25 | .806 | 0.124 ± 0.005 | 43.81 | 24.07 | < .**.001** |
| Nb | -0.149 ± 0.025 | 39.94 | -6.04 | < .**.001** | 0.264 ± 0.008 | 27.72 | 34.93 | < .**.001** |
| Pb | -0.222 ± 0.032 | 28.78 | -6.88 | < .**.001** | 0.370 ± 0.007 | 29.41 | 52.09 | < .**.001** |
| V | 0.030 ± 0.035 | 26.42 | 0.88 | .387 | -0.373 ± 0.007 | 27.70 | -51.19 | < .**.001** |
| Rate:Frequency | 0.025 ± 0.009 | 4099.49 | 2.83 | **.005** | -0.007 ± 0.002 | 5121.60 | -4.47 | < .**.001** |
| Rate:Pa | 0.027 ± 0.009 | 4091.66 | 2.92 | **.003** | -0.008 ± 0.002 | 5104.73 | -4.01 | < .**.001** |
| Rate:Nb | 0.045 ± 0.010 | 4105.64 | 4.70 | < .**.001** | -0.016 ± 0.002 | 5127.63 | -7.72 | < .**.001** |
| Rate:Pb | 0.084 ± 0.010 | 4110.07 | 8.58 | < .**.001** | -0.008 ± 0.002 | 5109.02 | -3.73 | < .**.001** |
| Rate:V | 0.054 ± 0.009 | 4090.21 | 5.86 | < .**.001** | 0.016 ± 0.002 | 5104.14 | 8.11 | < .**.001** |
| Frequency:Pa | 0.027 ± 0.019 | 4092.58 | 1.44 | .151 | 0.038 ± 0.003 | 5107.27 | 11.64 | < .**.001** |
| Frequency:Nb | -0.074 ± 0.020 | 4099.19 | -3.70 | < .**.001** | 0.101 ± 0.003 | 5128.21 | 29.35 | < .**.001** |
| Frequency:Pb | -0.147 ± 0.021 | 4113.15 | -7.02 | < .**.001** | 0.104 ± 0.004 | 5132.55 | 29.45 | < .**.001** |
| Frequency:V | 0.390 ± 0.019 | 4090.55 | 21.04 | < .**.001** | -0.070 ± 0.003 | 5104.78 | -21.72 | < .**.001** |

**Model 5:** $\log_2(\text{latency or amplitude}) \sim \log_2(\text{rate}) + \log_2(\text{frequency}) + \text{peak} + \log_2(\text{rate}): \log_2(\text{frequency}) + \log_2(\text{rate}): \text{peak} + \log_2(\text{frequency}): \text{peak} + (\log_2(\text{rate}) + \log_2(\text{frequency}) + \text{peak} + \text{ear} \mid \text{participant})$
Note: significant p-values are bolded.

Consistent with peak presence at 81 dB peSPL, wave V, Na, and Pa had the largest amplitudes (Fig. 3b; ANOVA of the mixed effects model, peak main effect *F*(4, 43.42) = 25.92, *p* < 0.001). The mean ± SE peak amplitudes across frequencies and rates were 0.14 ± 0.002 µV for wave V, 0.10 ± 0.003 µV for Na, 0.11 ± 0.003 µV for Pa, and 0.08 ± 0.002 µV for both Nb and Pb. Peak amplitudes also differed by tone-pip frequency (2-way interaction, *F*(4, 4099.24) = 225.88, *p* < 0.001). Whereas Na and Pa amplitudes were more similar between 500 and 8000 Hz (mixed model slope for frequency for Na, *p* = 0.452; change in slope for Pa, *p* = 0.150), wave V increased with higher frequencies (positive change in slope re: Na, *p* < 0.001) but Nb and Pb decreased (negative change in slope re: Na, *p* < 0.001). Thus, from earlier waves to later waves, the amplitudes went from increasing with frequency, to no change, to decreasing with frequency. Overall, peak amplitudes decreased with increasing stimulation rate (rate main effect, *F*(1, 29.25) = 2816.62, *p* < 0.001). But as a result of the different MLR patterns across frequency, the amplitude changes with rate were smaller/shallower for the higher frequencies (rate x frequency interaction, *F*(1, 4099.53) = 8.03, *p* = 0.005; positive, or less negative, change in slope) and for all other peaks compared to Na (2-way interaction, *F*(4, 4102.72 = 20.77, *p* < 0.001; all changes in slopes re: Na *p* < 0.003).

Peak latencies are shown in Fig. 3c and the model given in Table 1. As expected, Pa, Nb, and Pb occurred progressively later than Na but wave V occurred earlier (ANOVA of the mixed effects model, main effect of peak *F*(4, 40.37) = 4389.03, *p* < 0.001). Mean ± SE latency across frequencies and rates was 8.94 ± 0.07 ms for V, 20.87 ± 0.12 ms for Na, 27.52 ± 0.12 ms for Pa, 38.11 ± 0.15 ms for Nb, and 50.15 ± 0.17 ms for Pb. Latencies were also earlier for higher frequencies (main effect, *F*(1, 28.71) = 550.18, *p* < 001), especially for Na and wave V (peak x frequency interaction, *F*(4, 5109.46) = 901.63, *p* < 0.001; change in slope for V re: Na *p* < 0.001). However, latencies became more similar across tone-pip frequency for the later MLR peaks (positive changes in slopes to become less negative re: Na; *p* < 0.001 for Pa, Nb and Pb). While the overall main effect of rate was not significant (*F*(1, 26.00) = 1.13, *p* = 0.298), the MLR also behaved differently to wave V with increasing stimulation rate (peak x frequency interaction, *F*(4, 5109.46) = 70.85, *p* < 0.001). Compared to Na (slope *p* = 0.240), latency increased with increasing rate for wave V (positive change in slope, *p* < 0.001) but decreased for the other MLR waves (negative change in slopes, all *p* < 0.001). The changes in latency with rate were also slightly greater for higher frequencies (rate x frequency interaction, *F*(1, 5121.57) = 20.00, *p* < 0.001). Although these were significant effects, the latency changes for MLR were relatively minimal relative to their absolute latency (the maximum change in latency between 20 and 120 Hz was −3.8 ms for Nb, or 9.6% of the 20 Hz mean latency of 39.9 ms). Thus, normative values for latencies are given in Table 2 for each tone-pip frequency at the most used rate of 40 Hz.

**Table 2.** Normative latency values ± SD (ms) for 81 & 51 dB peSPL at 40 Hz rate.

|  | 500 Hz | 1000 Hz | 2000 Hz | 4000 Hz | 8000 Hz |
| --- | --- | --- | --- | --- | --- |
| **81 dB peSPL** |  |  |  |  |  |
| **V** | 12.7 ± 1.2 | 9.5 ± 0.9 | 8.0 ± 0.4 | 7.1 ± 0.3 | 6.7 ± 0.3 |
| **Na** | 26.7 ± 1.3 | 22.5 ± 1.7 | 20.0 ± 2.2 | 18.2 ± 2.3 | 17.0 ± 1.8 |
| **Pa** | 33.8 ± 1.4 | 29.7 ± 1.6 | 27.5 ± 1.7 | 25.3 ± 2.3 | 24.4 ± 2.7 |
| **Nb** | 43.0 ± 4.5 | 38.5 ± 2.5 | 38.6 ± 2.7 | 36.9 ± 3.4 | 36.3 ± 3.0 |
| **Pb** | 56.5 ± 2.1 | 51.1 ± 2.9 | 50.1 ± 3.6 | 47.9 ± 4.3 | 47.1 ± 4.3 |
| **51 dB peSPL** |  |  |  |  |  |
| **V** | 14.7 ± 1.0 | 11.6 ± 0.6 | 9.2 ± 0.3 | 8.1 ± 0.3 | 7.4 ± 0.4 |
| **Na** | 28.1 ± 2.1 | 23.9 ± 2.3 | 20.9 ± 2.1 | 19.0 ± 2.6 | 17.0 ± 2.3 |
| **Pa** | 35.3 ± 1.5 | 31.5 ± 1.3 | 28.9 ± 1.1 | 26.2 ± 2.4 | 24.9 ± 2.4 |
| **Nb** | 46.1 ± 2.8 | 40.9 ± 2.9 | 39.3 ± 2.0 | 37.6 ± 3.3 | 36.6 ± 2.3 |
| **Pb** | 57.3 ± 6.5 | 50.6 ± 5.3 | 50.4 ± 5.6 | 47.2 ± 5.2 | 47.1 ± 4.7 |

### ABR wave V and MLR waves Na and Pa remain present at a level near threshold

Waveforms were also recorded at 51 dB peSPL (∼33 dB nHL), a level closer to hearing threshold. Fig. 4a presents the grand average waveforms for both intensities at 40 Hz, which is the single optimal stimulation rate previously identified for the pABR and commonly used in clinical practice to estimate hearing thresholds [14]. Data for all rates are included in Online Resource 2 and the normative latencies for 40 Hz are given in Table 2. At the near-threshold lower level, ABR wave V and MLR components Na and Pa remained robust (Fig. 4a) and maintained high detectability across frequency (Fig. 4b, V: 100%, Na: 96%, Nb: 96%). Similarly, Nb and Pb maintained detectability of ∼87% on average, but P0 was less identifiable at this lower level (65%) and was therefore excluded from analysis. Unsurprisingly, ABR waves I and III were rarely present at this near-threshold level, averaging 27.5% and 28.5%, respectively, and were therefore also excluded from subsequent analysis.

**Fig. 4.**
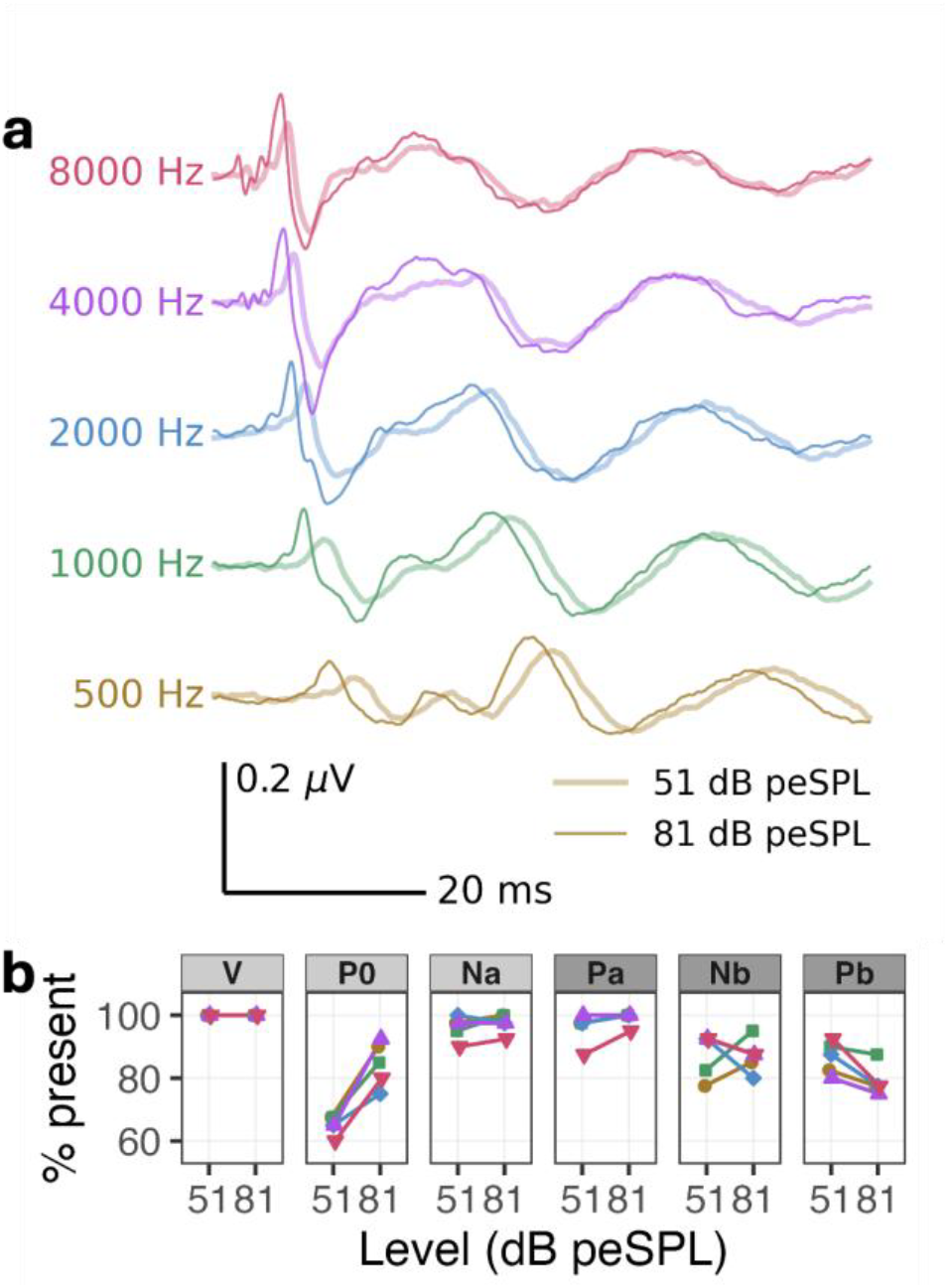
(a) Grand average responses at 81 dB peSPL (dark thin line) and 51 dB peSPL (light thick line) levels at the 40 Hz stimulation rate. Subject-ears are averaged for display purposes. (b) Wave presence for 20 participants for both levels

Changes in amplitude and latency between the two levels were quantified using the ratio of 51 dB peSPL to 81 dB peSPL. The linear mixed effects model is provided in Table 3. Fig. 5 illustrates the effects of decreasing the stimulus level as density plots with horizontal error bars indicating the 95% confidence interval of the mean ratio. The vertical gray line marks a ratio of 1, corresponding to no change between stimulus levels. Rightward distributions indicate an increase in ratio (larger or later peak), while leftward distributions indicate a decrease in ratio (smaller or earlier peak).

**Table 3.** Linear mixed effect models for 51 dB peSPL.

|  | Amplitude |  |  |  | Latency |  |  |  |
| --- | --- | --- | --- | --- | --- | --- | --- | --- |
|  | *Estimate ± SE* | *df* | *t* | *p* | *Estimate ± SE* | *df* | *t* | *p* |
| Intercept (Na) | 0.898 ± 0.022 | 76.62 | 41.72 | < .001 | 1.059 ± 0.007 | 121.44 | 141.41 | < .001 |
| Nb | 0.067 ± 0.027 | 488.50 | 2.52 | .012 | -0.012 ± 0.010 | 528.23 | -1.19 | .233 |
| V | -0.154 ± 0.024 | 481.03 | -6.50 | < .001 | 0.115 ± 0.009 | 523.14 | 12.71 | < .001 |
| Frequency | 0.003 ± 0.024 | 482.95 | 0.11 | .913 | -0.046 ± 0.009 | 526.02 | -5.07 | < .001 |

**Model 2:** ratio ~ peak + log<sub>10</sub>(frequency) + (1 | participant)

**Fig. 5.**
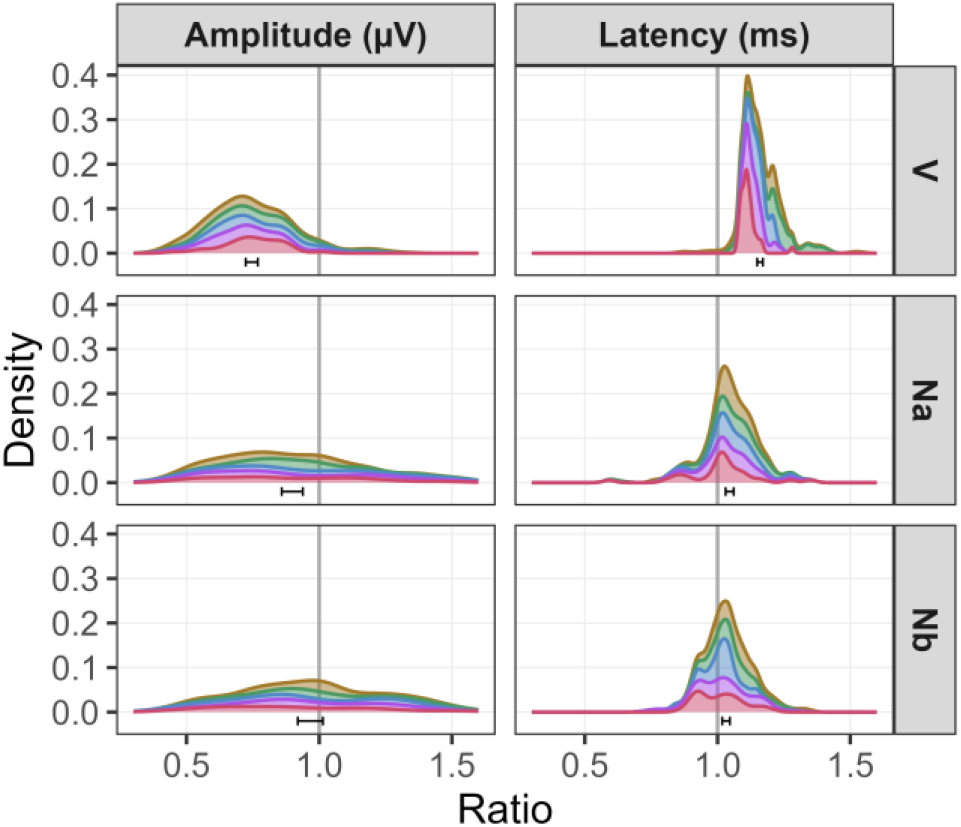
Density plots of amplitude and latency ratios, calculated as 51 dB peSPL divided by 81 dB peSPL, for identified peaks. Colors indicate stimulus frequency. Horizontal error bars represent the 95% confidence interval of the mean ratio from a two-tailed t-test

Overall, amplitude ratios were similar across the tone-pip frequencies (AVOVA of the mixed effects model: *F*(1, 482.95) = 0.012, *p* = 0.913) but differed by peak (*F*(2, 485.82), *p* < 0.001). Specifically, Na amplitude decreased to a ratio of 0.90 (95% confidence interval, 0.86–0.94) and wave V decreased further to 0.75 (0.72–0.77; change in intercept from Na, *p* < 0.001). Both ratios represented significant amplitude reductions (independent *t*- tests with µ=1, FDR-corrected *p* < 0.001; V: *t*(197) = −22.00; Na: *t*(176) = −5.04). In contrast, Nb amplitude ratios were more variable, resulting in a smaller overall change in amplitude to a ratio of 0.97 (0.919–1.014; *p* = 0.233) that did not represent a significant amplitude change when level was lowered (independent *t*-test, *t*(127) = −1.398, FDR-corrected *p* = 0.165). However, Nb amplitude was smaller at the 81 dB peSPL level to begin with (Fig. 3b), likely contributing to the variable range of amplitude ratios seen in Fig. 5a.

Unlike peak amplitude, all peak latencies increased overall (i.e., mean ratios (95% confidence intervals) > 1, independent t-tests with µ=1, all FDR-corrected p < 0.001; V: 1.13 (1.15–1.17), t(199) = 29.5; Na: 1.05 (1.03– 1.06), t(186) = 5.9; Nb: 1.03 (1.02–1.05), t(156) = 4.4) and the latency delay was greater for lower-than higher-frequency responses (mixed effects model, p < 0.001). Like peak amplitude ratios, the latency ratios reflected a greater delay (i.e., bigger difference from 1) for wave V than Na (mixed model change in intercept for V re: Na, p < 0.001), but a similar delay for the two MLR peaks (mixed model change in intercept for Nb re: Na, p = 0.233). Thus, these ratios indicated the MLR waves changed less than wave V with a level change.

## Discussion

This study aimed to evaluate whether MLR components could be reliably recorded alongside the ABR using pABR parameters optimized for wave V detection, and to characterize MLR responses for tone-pip frequencies across changes in recording parameters. Most participants had clearly visible ABR wave V and MLR components Na-Pa and Nb-Pb (96–100% and 79–88% respectively) at both a moderate suprathreshold level (81 dB peSPL, ∼63 dB nHL) and a lower, near-threshold level (51 dB peSPL, ∼33 dB nHL). Compared to ABR, MLR peaks remained more similar across changes in rate, frequency and level. This was particularly evident for a decrease in level, where MLR waves became just as, if not more, prominent than wave V. Thus, despite the faster stimulation rates, consistently visible/present Na-Pa and Nb-Pb complexes suggest simultaneous recording of ABR and MLR is feasible at soft and moderate levels. This supports the potential clinical value of viewing both responses together to aid response interpretation, particularly when wave V is smaller and MLR components more prominent.

### Simultaneous ABR and MLR recording is feasible under ABR-optimized conditions

With pABR-based parameters and recording conditions, MLRs were consistently identifiable. In fact, the Na-Pa components were present in at least 98% of participants at 81 dB peSPL and 96% of cases at 51 dB peSPL, despite relaxed/sleeping arousal states and suboptimal MLR recording parameters [8, 28]. Furthermore, at the lower level, Na-Pa amplitudes did not change as much as wave V (Fig 5) and even exceeded those of wave V across all frequencies and rates (Fig 4), consistent with prior reports that Na-Pa components are less affected by intensity changes than wave V [8, 21, 23, 28]. This robustness of Na-Pa may arise from synchronous firing across the multiple overlapping thalamo-cortical inputs [7, 8, 21, 28], which may also contribute to their broader morphology. Overall, these high detection rates and robust amplitudes show that MLRs can be recorded simultaneously with ABRs and may serve as a useful complement for assessing sound processing at low to moderate levels.

Although wave V and Na-Pa were robustly measured in the simultaneous recordings, other ABR and MLR components were less prominent and harder to identify. Consistent with long-standing previous literature [2, 4, 15, 26], wave I and III detection rates declined substantially with faster rates and the lower intensity (Fig 3). Even at the louder suprathreshold level, only 36–56% of waves I and III were present, suggesting that a moderate level is not sufficient to evaluate early wave metrics. However, there may be one exception: this study included 8000 Hz, and the moderate level elicited waves I and III in 85–97.5% of participants. These high-frequency ABRs at a moderate level might be worth exploring given the links between high frequency hearing and speech-in-noise abilities [32– 36]. Thus, while earlier waves may be absent at lower frequencies, high frequency waves could still provide valuable information at the more moderate levels without adding to recording time.

MLR components P0, Nb, and Pb were also less consistently present than Na-Pa across participants (Fig. 3). The earliest MLR wave, P0, was the least detectable MLR wave, consistent with reports across studies [6, 22, 23, 25, 28]. The extended analysis windows allowed us to confirm postauricular muscle responses (PAM) were not present and did not affect P0 identification in 18 of the 20 participants (see Online Resource 4 for responses from the 2 participants that exhibited a PAM). More likely, the resting /lower arousal levels of the participants contributed to this variable presence across the 18 participants without a PAM [8, 21, 27–29]. The primary neural generator of P0 is the inferior colliculus, which is involved with integrating both arousal related inputs and auditory signals [7, 8, 25, 28]. The later MLR waves Nb and Pb arise mainly from the primary and association cortices, also making them vulnerable to state-dependent modulation and suppression. Sleep is known to diminish or abolish these components [7, 8, 24, 28]. For this study, rest/sleep was encouraged to optimize pABR recording [14] which could have reduced the amplitude and visibility of these MLR waves. While this methodological choice may limit visibility of later components, it is the standard practice in many clinical protocols. Nb and Pb might be more visible under different (i.e., more awake) arousal states. However, the numbers reported herein provide realistic insight into how these responses would occur if MLR was added to the clinical protocol with simultaneous recording paradigms.

### MLR amplitude and latency remained more consistent than wave V across parameters changes

MLR components changed across parameters to a lesser extent than wave V. At the suprathreshold level, Na and Pa were the largest MLR peaks (both ∼0.11 µV across rates and frequencies), second only to wave V (∼0.15 µV). But at the lower level, Na-Pa amplitude did not change as much as wave V, becoming similarly sized (∼0.09 versus 0.11 µV respectively). Frequency effects on Na and Pa were modest: their amplitudes did not differ significantly across tone-pip frequencies, in contrast to the larger wave V amplitudes at higher frequencies. In addition, while wave V showed the well-known latency increase with increasing rate [2, 4, 15, 26], MLR components showed the opposite: with progressively later components, the latencies became slightly *earlier*. This latency pattern aligns with previous studies on MLR responses to click stimuli [8, 23, 27, 28] and could reflect aspects pertaining to the broader morphology of the MLR components. First, while statistically significant, the latency shifts were relatively small (∼3.5 ms across rates for Na-Pa and Nb-Pb) in the context of their broader, later latency range (i.e., ∼10% of their absolute latencies overall). Second, these latency shifts could reflect the less precise nature of picking a peak from a broader wave compared to the sharper wave V. Third, overlapping neural generators may impact the exact peak timing of the potential measured on the surface of the head. Taken together, these results indicate that unlike wave V, MLR components are less affected by tone-pip frequency, and their latency ranges remain essentially constant across the higher stimulation rates used in ABR protocols.

This study provides new normative data on adult MLR components across tone-pip frequencies and a higher range of stimulus rates that would be used in a simultaneous pABR/MLR paradigm. Given the similarity of latency across rates, the normative values are given for the optimal ABR rate of 40 Hz (Table 2). The ±1SD ranges (68% of the data) of our tone-pip data for Na and Pa mostly overlap with the ranges provided in previous studies that used comparable levels (50 dB SL or 60–70 dB nHL), both a low- and high-frequency stimulus (500/1000 Hz and 3000/4000 Hz or click), but slower 8-12 Hz rates [7, 17, 20, 21, 23]. The overlap in latency ranges, despite differing rates, is consistent with the very minimal change in latency seen with increasing rate (Fig. 3). The only difference from previous work is that our low-frequency Na peaks were later (500 Hz: 25–28ms compared to 20–22 ms) [21]. The later latencies could be due to the higher prevalence of P0 identified in our lower-frequency MLRs than previously reported (ours: >78%, previous: <50-60% or not described due to low incidence) [6, 21–23, 25, 28]. The overall similar values across different stimulation rates suggest that simultaneously recording MLR with ABR does not significantly impact where a clinician would look to identify these component peaks.

### Na-Pa components provide additional context for clinical interpretation

Although Na-Pa amplitudes decrease slightly at higher rates, they remain prominent with minimal latency changes at both levels. These characteristics make the Na-Pa complex a strong candidate for providing complementary diagnostic information when included alongside wave V. At 81 dB peSPL (∼63 dB nHL), reliable Na-Pa responses were obtained, capturing a broader portion of the auditory system without added test time. This moderate level better reflects real-world listening and improves comfort compared to traditionally higher suprathreshold intensities [32, 33, 35, 36]. At lower levels (51 dB peSPL, ∼33 dB peSPL), the MLR provides additional evidence for whether a response is present and makes it easier to identify broader low-frequency wave Vs that are harder to distinguish from recording noise. Fig. 6 demonstrates this potential diagnostic value by showing one participant’s 500 Hz response at 51 dB peSPL and 40 Hz, parameters typical in clinical protocols for threshold estimation. In the ABR-typical 20 ms analysis window (Fig. 6, top waveform), wave V appears broad and less distinct, making it harder to determine if there is a reliable response, and thus, if the level is above threshold. Expanding the view to include the MLR (Fig. 6b) reveals prominent Na-Pa and Nb-Pb waves, confirming auditory pathway activation and supporting a more confident judgement that the wave V is present, and the level is above threshold. This added evidence may improve accuracy of threshold predictions by enabling more confident choices of lower levels, especially for low-frequency tone-pips, which are known to yield the least accurate threshold estimates [10, 17–19]. While we did not directly evaluate threshold estimation, our results suggest that incorporating Na-Pa into simultaneous recordings may aid wave V detection and support more confident peak identification for this more challenging frequency response.

**Fig. 6.**
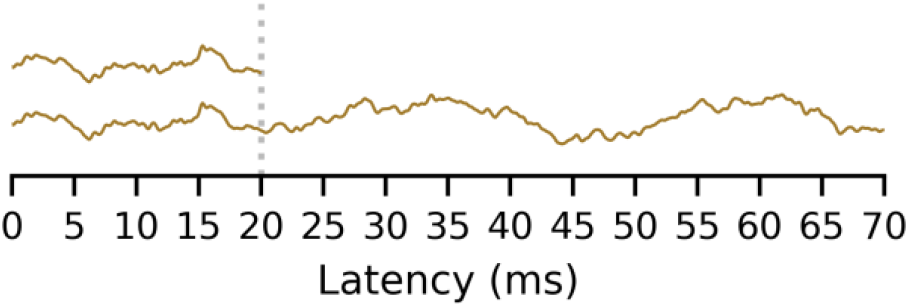
Example of an individual participant’s 500 Hz waveform recorded at 51 dB peSPL with a 40 Hz stimulation rate. Wave V is broader and smaller (top) and less distinguishable from the background noise, but the MLR Na-Pa (bottom) supports its presence as an indication of near/above threshold

## Limitations and Future Directions

Despite strong evidence for simultaneous ABR-MLR recording, targeted follow up studies are needed. This work comprised a secondary analysis of an existing dataset [1] that was not specifically designed for MLR recordings. Participants were encouraged to rest, sleep, or watch closed-captioned videos, meaning that their likely low-arousal state may have reduced visibility of later cortical components such as Nb and Pb, which are sensitive to participant state [7, 8, 24, 28]. Similarly, testing infants is a particularly important next step, as MLR generators are still developing in early childhood and could be more susceptible to arousal state, and later components such as Nb-Pb are less consistently observed at young ages [8, 24, 27, 29]. Finally, we did not assess responses at behavioral threshold or across a broad range of intensities. Future studies could evaluate these systematic MLR changes with level and the accuracy of predicting thresholds when considering the MLR’s presence. Together, these factors outline future opportunities to broaden and improve ABR-MLR testing across different populations, protocols, and clinical contexts.

## Conclusion

This study demonstrates that simultaneously recording ABR and MLR responses is feasible using the pABR paradigm with parameters optimized for ABR. Across a range of stimulation rates and frequencies, MLR components Na and Pa were consistently observed, even at near threshold levels, whereas waves I, III and other MLR peaks (P0, Nb, Pb) were less reliably detected. Na-Pa and Nb-Pb retained their amplitude at the lower level, suggesting they may aid wave V detection, especially for low frequency responses with broader and harder to identify waveforms closer to threshold. This added information could enhance the accuracy and confidence of threshold estimation, especially in clinical contexts where precise identification of hearing thresholds at specific frequencies is critical. Future research should extend this work to infants and individuals with hearing loss and further explore MLR visibility at behavioral thresholds. Overall, these results provide a strong foundation for integrated evoked response testing and highlight the potential of the pABR to broaden clinical and research applications in auditory assessment.

## Supporting information

Online Resources

## Author Contribution Statement

Conceptualization: Melissa Polonenko; Data Curation: Isabel Herb; Formal Analysis and Investigation: Isabel Herb, Melissa Polonenko; Methodology: Isabel Herb, Melissa Polonenko; Resources: Melissa Polonenko; Supervision: Melissa Polonenko; Visualization: Isabel Herb, Melissa Polonenko, Writing – Original Draft: Isabel Herb; Writing – Review & Editing: Isabel Herb, Melissa Polonenko.

## Statements and Declarations

### Data Availability Statement

The electroencephalography responses were downloaded from an openly available dataset [1]. Additional data that support the findings of this study and the associated analysis code will be available on GitHub at https://github.com/polonenkolab/pabr_simultaneous_abrmlr.,

## Acknowledgements

This work was supported by the University of Minnesota Department of Speech-Language-Hearing Sciences Summer Fellowship.

## References

1. Polonenko M, Maddox R (2021) Optimizing parameters for using the parallel auditory brainstem response (pABR) to quickly estimate hearing thresholds. Dryad. 10.5061/dryad.1c59zw3vm

2. Don M, Eggermont J (1978) Analysis of the click-evoked brainstem potentials in man unsing high-pass noise masking. J Acoust Soc Am 63:1084–1092. 10.1121/1.381816

3. Hecox K, Galambos R (1974) Brain stem auditory evoked responses in human infants and adults. Arch Otolaryngol Chic Ill 1960 99:30–33. 10.1001/archotol.1974.00780030034006

4. Jewett D, Williston J (1971) Auditory-evoked far fields averaged from the scalp of humans. Brain J Neurol 94:681–696. 10.1093/brain/94.4.681

5. Møller AR, Jannetta PJ (1983) Interpretation of brainstem auditory evoked potentials: results from intracranial recordings in humans. Scand Audiol 12:125–133. 10.3109/01050398309076235

6. Picton TW, Hillyard SA, Krausz HI, Galambos R (1974) Human auditory evoked potentials. I. Evaluation of components. Electroencephalogr Clin Neurophysiol 36:179–190. 10.1016/0013-4694(74)90155-2

7. Liégeois-Chauvel C, Musolino A, Badier JM, et al (1994) Evoked potentials recorded from the auditory cortex in man: evaluation and topography of the middle latency components. Electroencephalogr Clin Neurophysiol 92:204–214. 10.1016/0168-5597(94)90064-7

8. McGee T, Kraus N (1996) Auditory development reflected by middle latency response. Ear Hear 17:419–429. 10.1097/00003446-199610000-00008

9. Musiek F, Geurkink N (1981) Auditory brainstem and middle latency evoked response sensitivity near threshold. Ann Otol Rhinol Laryngol 90:236–240. 10.1177/000348948109000308

10. Gorga MP, Johnson TA, Kaminski JR, et al (2006) Using a Combination of Click- and Tone Burst–Evoked Auditory Brain Stem Response Measurements to Estimate Pure-Tone Thresholds. Ear Hear 27:60–74. 10.1097/01.aud.0000194511.14740.9c

11. Stapells DR, Oates P (1997) Estimation of the pure-tone audiogram by the auditory brainstem response: a review. Audiol Neurootol 2:257–280. 10.1159/000259252

12. Stapells DR, Picton TW (1981) Technical aspects of brainstem evoked potential audiometry using tones. Ear Hear 2:20–29. 10.1097/00003446-198101000-00006

13. Polonenko M, Maddox R (2019) The Parallel Auditory Brainstem Response. Trends Hear 23:233121651987139. 10.1177/2331216519871395

14. Polonenko M, Maddox R (2022) Optimizing Parameters for Using the Parallel Auditory Brainstem Response to Quickly Estimate Hearing Thresholds. Ear Hear 43:646–658. 10.1097/AUD.0000000000001128

15. Valderrama JT, Alvarez I, de la Torre A, et al (2012) Recording of auditory brainstem response at high stimulation rates using randomized stimulation and averaging. J Acoust Soc Am 132:3856–3865. 10.1121/1.4764511

16. Musiek F, Baran J (2022) Neuroaudiological Considerations for the Auditory Brainstem Response and Middle Latency Response Revisited: Back to the Future. Semin Hear 43:149–161. 10.1055/s-0042-1756161

17. Hayes D, Jerger J (1982) Auditory brainstem response (ABR) to tone-pips: results in normal and hearing-impaired subjects. Scand Audiol 11:133–142. 10.3109/01050398209076210

18. Kileny P (1981) The Frequency Specificity of Tone-Pip Evoked Auditory Brain Stem Responses: Ear Hear 2:270–275. 10.1097/00003446-198111000-00006

19. Stapells DR, Gravel JS, Martin BA (1995) Thresholds for auditory brain stem responses to tones in notched noise from infants and young children with normal hearing or sensorineural hearing loss. Ear Hear 16:361– 371. 10.1097/00003446-199508000-00003

20. Barajas JJ, Exposito M, Fernandez R, Martin LJ (1988) Middle latency response to a 500-Hz tone pip in normal-hearing and in hearing-impaired subjects. Scand Audiol 17:21–26. 10.3109/01050398809042176

21. Maurizi M, Ottaviani F, Paludetti G, et al (1984) Middle-latency auditory components in response to clicks and low- and middle-frequency tone pips (0.5-1 kHz). Audiol Off Organ Int Soc Audiol 23:569–580. 10.3109/00206098409081539

22. McFarland WH, Vivion MC, Goldstein R (1977) Middle components of the AER to tone-pips in normal-hearing and hearing-impaired subjects. J Speech Hear Res 20:781–798. 10.1044/jshr.2004.781

23. Thornton AR, Mendel MI, Anderson CV (1977) Effects of stimulus frequency and intensity on the middle componenets of the averaged auditory eletroencephalic response. J Speech Hear Res 20:81–94. 10.1044/jshr.2001.81

24. Musiek F, Nagle S (2018) The Middle Latency Response: A Review of Findings in Various Central Nervous System Lesions. J Am Acad Audiol 29:855–867. 10.3766/jaaa.16141

25. Hashimoto I (1982) Auditory evoked potentials from the human midbrain: slow brain stem responses. Electroencephalogr Clin Neurophysiol 53:652–657. 10.1016/0013-4694(82)90141-9

26. Holt F, Özdamar Ö (2016) Effects of rate (0.3-40/s) on simultaneously recorded auditory brainstem, middle and late responses using deconvolution. Clin Neurophysiol Off J Int Fed Clin Neurophysiol 127:1589–1602. 10.1016/j.clinph.2015.10.046

27. Jerger J, Chmiel R, Glaze D, Frost JD (1987) Rate and filter dependence of the middle-latency response in infants. Audiol Off Organ Int Soc Audiol 26:269–283. 10.3109/00206098709081555

28. Ozdamar O, Kraus N (1983) Auditory middle-latency responses in humans. Audiol Off Organ Int Soc Audiol 22:34–49. 10.3109/00206098309072768

29. Okitsu T (1984) Middle components of the auditory evoked response in young children. Scand Audiol 13:83–86. 10.3109/01050398409043044

30. Ontario Ministry of Children, Community, and Social Services (2019) Ontario Infant Hearing Program: Protocol for Universal Hearing Screening in Ontario

31. The Joint Committee on Infant Hearing (2019) Year 2019 Position Statement: Principles and Guidelines for Early Hearing Detection and Intervention Programs. JEHDI. 10.15142/FPTK-B748

32. Bharadwaj HM, Mai AR, Simpson JM, et al (2019) Non-Invasive Assays of Cochlear Synaptopathy – Candidates and Considerations. Neuroscience 407:53–66. 10.1016/j.neuroscience.2019.02.031

33. Bramhall N, Beach EF, Epp B, et al (2019) The search for noise-induced cochlear synaptopathy in humans: Mission impossible? Hear Res 377:88–103. 10.1016/j.heares.2019.02.016

34. Mehraei G, Hickox AE, Bharadwaj HM, et al (2016) Auditory Brainstem Response Latency in Noise as a Marker of Cochlear Synaptopathy. J Neurosci 36:3755–3764. 10.1523/JNEUROSCI.4460-15.2016

35. Plack CJ, Barker D, Prendergast G (2014) Perceptual Consequences of “Hidden” Hearing Loss. Trends Hear 18:2331216514550621. 10.1177/2331216514550621

36. Prendergast G, Tu W, Guest H, et al (2018) Supra-threshold auditory brainstem response amplitudes in humans: Test-retest reliability, electrode montage and noise exposure. Hear Res 364:38–47. 10.1016/j.heares.2018.04.002

37. Dallos P (1992) The active cochlea. J Neurosci 12:4575–4585. 10.1523/JNEUROSCI.12-12-04575.1992

38. Maddox RK (2020) S/Plitter: Hardware and firmware for converting digital audio to TTL triggers (version 2) [computer program]

39. Elberling C, Wahlgreen O (1985) Estimation of auditory brainstem response, ABR, by means of Bayesian inference. Scand Audiol 14:89–96. 10.3109/01050398509045928

40. Koo TK, Li MY (2016) A Guideline of Selecting and Reporting Intraclass Correlation Coefficients for Reliability Research. J Chiropr Med 15:155–163. 10.1016/j.jcm.2016.02.012

41. Burnham KP, Anderson DR, Huyvaert KP (2011) AIC model selection and multimodel inference in behavioral ecology: some background, observations, and comparisons. Behav Ecol Sociobiol 65:23–35. 10.1007/s00265-010-1029-6

42. Benjamini Y, Hochberg Y (1995) Controlling the False Discovery Rate: A Practical and Powerful Approach to Multiple Testing. J R Stat Soc Ser B Stat Methodol 57:289–300. 10.1111/j.2517-6161.1995.tb02031.x

43. Posit team (2025) RStudio: Integrated Development Environment for R. Posit Software, PBC, Boston, MA

44. R Core Team (2025) R: A Language and Environment for Statistical Computing. R Foundation for Statistical Computing, Vienna, Austria

45. Revelle (2025) psych: Procedures for Psychological, Psychometric, and Personality Research. Northwestern University, Evanston, Illinois

46. Bates D, Mächler M, Bolker B, Walker S (2015) Fitting Linear Mixed-Effects Models Using lme4. J Stat Softw 67:. 10.18637/jss.v067.i01

47. Bolker B, Robinson D (2024) broom.mixed: Tidying Methods for Mixed Models

48. Kuznetsova A, Brockhoff PB, Christensen RHB (2017) lmerTest Package: Tests in Linear Mixed Effects Models. J Stat Softw 82:1–26. 10.18637/jss.v082.i13

49. Wickham H, Averick M, Bryan J, et al (2019) tidyverse. J Open Source Softw 4:1686. 10.21105/joss.01686

50. Gramfort A (2013) MEG and EEG data analysis with MNE-Python. Front Neurosci 7:. 10.3389/fnins.2013.00267

