## Supplementary material for "Characterizing Simultaneously Recorded Auditory Brainstem and Middle Latency Responses Using the Parallel Auditory Brainstem Response Paradigm": Online Resources

**JARO – Journal of the Association for Research in Otolaryngology**

**Online Resource 1** AICc Model Comparison for 81 dB peSPL

| | | Amplitude (no 8 kHz) | | | Latency | | |
|---|---|---|---|---|---|---|---|
| | *k* | *AICc* | *Δ AICc* | *AICc Wt.* | *AICc* | *Δ AICc* | *AICc Wt.* |
| **Model 5** | **53** | **-4482.82** | **0** | **0.92** | **-20535.65** | **5.42** | **0.06** |
| Model 4 | 57 | -4477.83 | 4.99 | 0.08 | -20541.07 | 0 | 0.94 |
| Model 3 | 22 | -3352.25 | 1130.57 | 0 | -19499.02 | 1042.05 | 0 |
| Model 2 | 9 | -2714.96 | 1767.85 | 0 | -17063.42 | 3477.65 | 0 |
| Model 1 | 3 | 269.69 | 4752.51 | 0 | 1283.19 | 21824.26 | 0 |

**Note:** Frequency is in kHz. Rate had 19 subtracted so 20 Hz was the intercept in the log-log model. The chosen model is bolded.

**Model 1:** $\log_{10}$(latency or amplitude) ~ (1 | participant)
**Model 2:** $\log_{10}$(latency or amplitude) ~ $\log_{10}$(rate) + $\log_{10}$(frequency) + peak + (1 | participant)
**Model 3:** $\log_{10}$(latency or amplitude) ~ $\log_{10}$(rate) + $\log_{10}$(frequency) + peak + $\log_{10}$(rate): $\log_{10}$(frequency) + $\log_{10}$(rate):peak + $\log_{10}$(frequency):peak + $\log_{10}$(rate): $\log_{10}$(frequency):peak + (1 | participant)
**Model 4:** $\log_{10}$(latency or amplitude) ~ $\log_{10}$(rate) + $\log_{10}$(frequency) + peak + $\log_{10}$(rate): $\log_{10}$(frequency) + $\log_{10}$(rate):peak + $\log_{10}$(frequency):peak + $\log_{10}$(rate): $\log_{10}$(frequency):peak + ($\log_{10}$(rate) + $\log_{10}$(frequency) + peak + ear | participant)
**Model 5:** $\log_{10}$(latency or amplitude) ~ $\log_{10}$(rate) + $\log_{10}$(frequency) + peak + $\log_{10}$(rate): $\log_{10}$(frequency) + $\log_{10}$(rate):peak + $\log_{10}$(frequency):peak + ($\log_{10}$(rate) + $\log_{10}$(frequency) + peak + ear | participant)

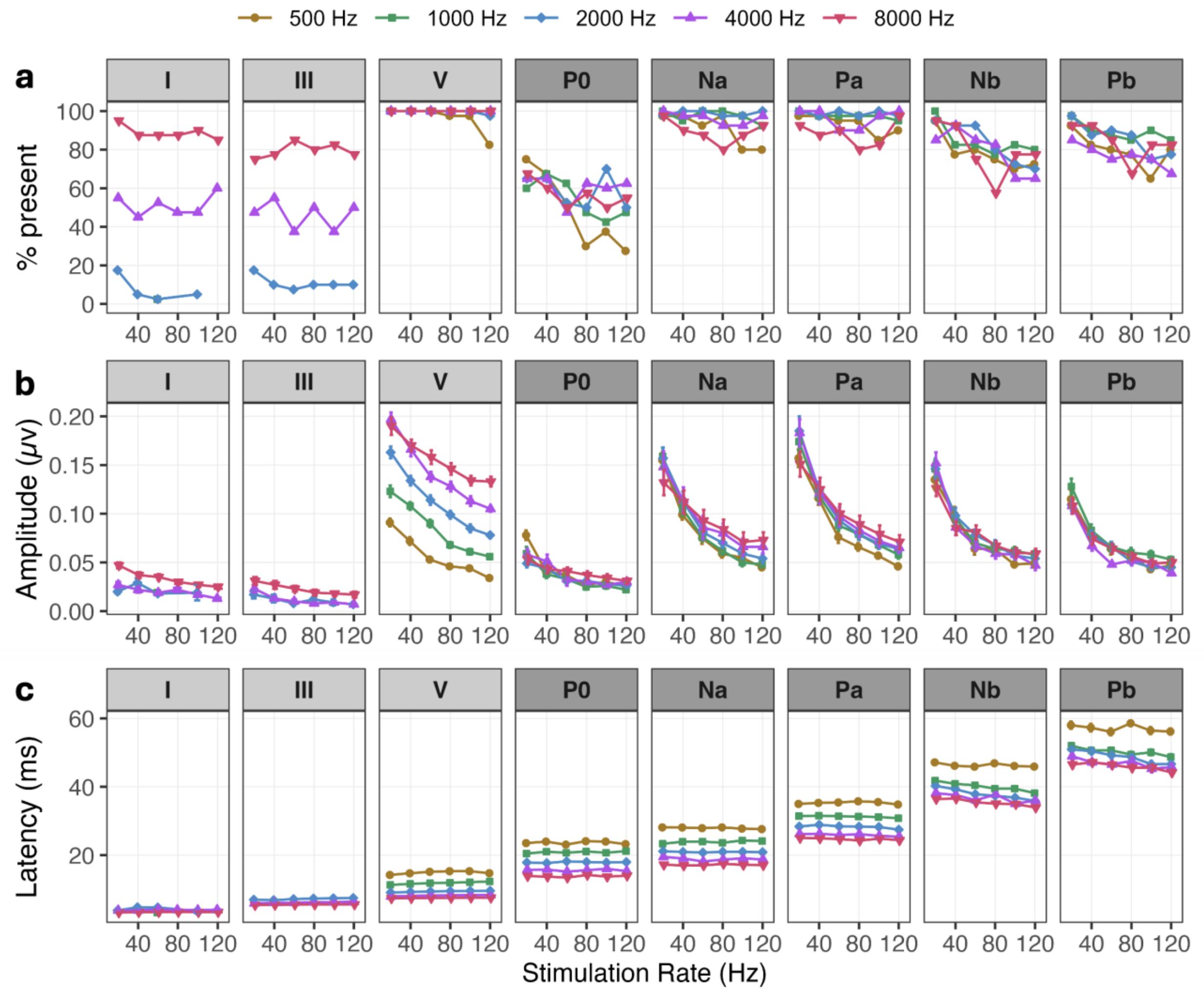


**Online Resource 2** Wave (a) presence across participant ears, as well as the mean (b) amplitude and (c) latency for stimuli presented at 51 dB peSPL across 6 rates and 5 tone-pip frequencies. Colors and shapes denote tone-pip frequency. Error bars (where large enough to be seen) indicate standard error of the mean. Panel shading delineates ABR (light gray) from MLR (dark gray) waves.

**Online Resource 3** AICc Model Comparison for 51 dB peSPL

| | | **Amplitude** | | | **Latency** | | |
|---|---|---|---|---|---|---|---|
| | *k* | *AICc* | *Δ AICc* | *AICc Wt.* | *AICc* | *Δ AICc* | *AICc Wt.* |
| Model 5 | 10 | -36.41 | 0 | 0.7 | -1072.62 | 0 | 0.66 |
| **Model 2** | **6** | **-33.98** | **2.43** | **0.21** | **-1070.76** | **1.86** | **0.26** |
| Model 3 | 8 | -32.31 | 4.10 | 0.09 | -1068.32 | 4.3 | 0.08 |
| Model 4 | 35 | 1.96 | 38.37 | 0 | -1038.57 | 34.05 | 0 |
| Model 1 | 3 | 36.18 | 72.59 | 0 | -864.65 | 207.97 | 0 |

**Note:** The chosen model is bolded.

Model 1: ratio ~ (1 | participant)
**Model 2: ratio ~ peak + log10(frequency) + (1 | participant)**
Model 3: ratio ~ peak * log10(frequency) + (1 | participant)
Model 4: ratio ~ peak * log10(frequency) + (1 + ear + peak * frequency | participant)
Model 5: Amplitude: ratio ~ peak + (peak | participant)
Latency: ratio ~ peak + log10(frequency) + (log10(frequency) | participant)

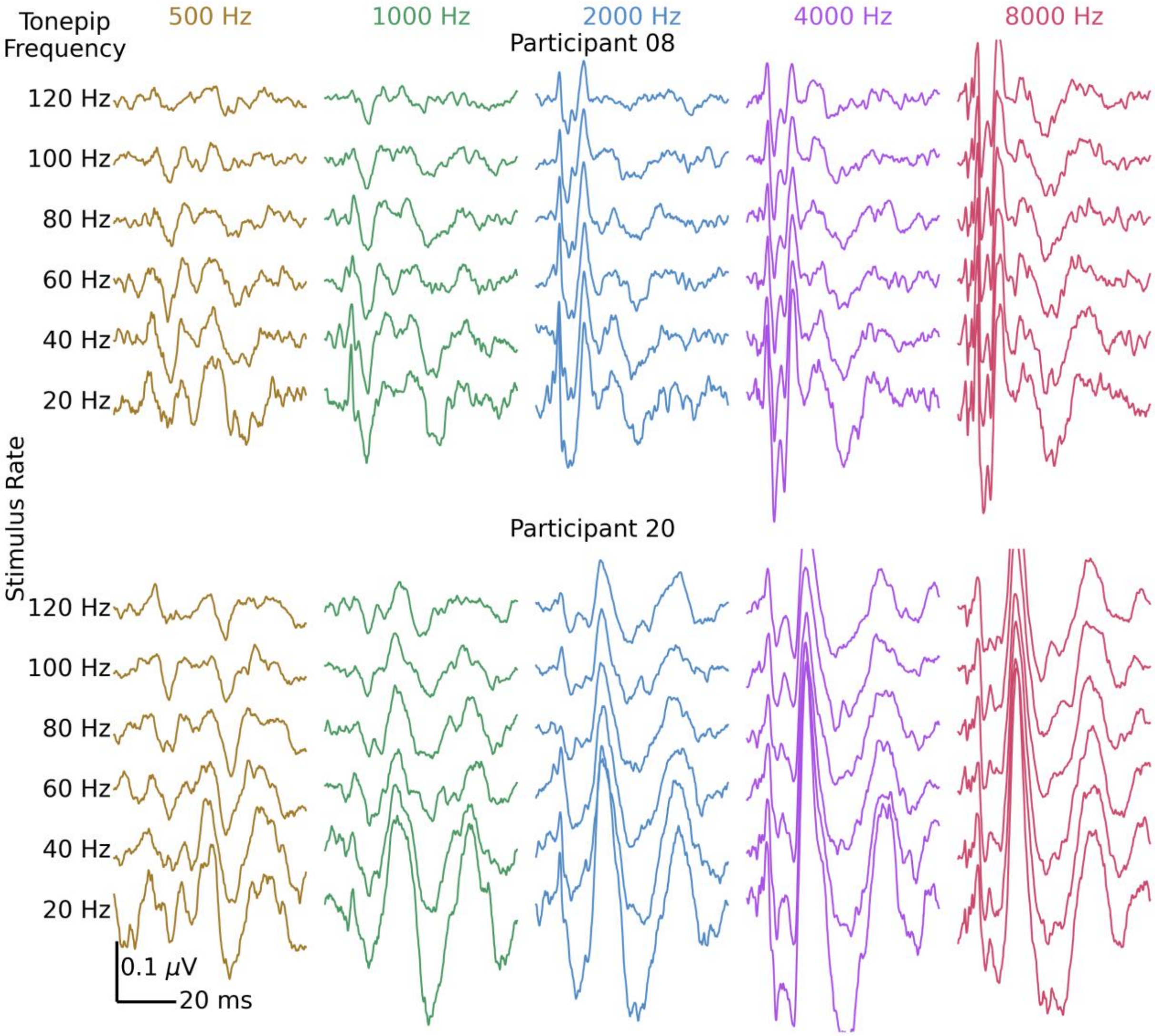


**Online Resource 4** Average waveforms for 2 participants at 81 dB peSPL across rates and frequencies. Both participants have a post-auricular muscle response(PAMR) interfering with early MLR peak components.
